# The nuclear to cytoplasmic ratio directly regulates zygotic transcription in *Drosophila* through multiple modalities

**DOI:** 10.1101/766881

**Authors:** Sahla Syed, Henry Wilky, João Raimundo, Bomyi Lim, Amanda A. Amodeo

**Author notes:** S.S. and H.W. contributed equally to this work.

## Abstract

Early embryos must rapidly generate large numbers of cells to form an organism. Many species accomplish this through a series of rapid, reductive, and transcriptionally silent cleavage divisions. Previous work has demonstrated that the number of divisions before both cell cycle elongation and zygotic genome activation (ZGA) is regulated by the ratio of nuclear content to cytoplasm (N/C). To understand how the N/C ratio affects the timing of ZGA, we directly assayed the behavior of several previously identified N/C-ratio-dependent genes using the MS2-MCP reporter system in living *Drosophila* embryos with altered ploidy and cell cycle durations. For every gene that we examined, we found that nascent RNA output per cycle is delayed in haploid embryos. Moreover, we found that the N/C ratio influences transcription through three separate modes of action. For some genes (*knirps* and *snail*) the effect of ploidy can be entirely accounted for by changes in cell cycle duration. However, for other genes (*giant, bottleneck and fruhstart*) the N/C ratio directly affects ZGA. For *giant* and *bottleneck,* the N/C ratio regulates the kinetics of transcription activation, while for *fruhstart* it controls the probability of transcription initiation. Our data demonstrate that the regulatory elements of N/C-ratio-dependent genes respond directly to the N/C ratio, through multiple modes of regulation, independent of interphase length.

## Introduction

The early embryo of many fast, externally developing species is largely transcriptionally silent during the rapid cleavage stage preceding a developmental transition known as the midblastula transition (MBT) (1, 2). At the MBT, the cell cycle slows and large-scale zygotic genome activation (ZGA) occurs (3–10). The timing of both cell cycle slowing and ZGA are controlled by the ratio of nuclear material, likely DNA, to cytoplasm (N/C ratio), as seen from previous studies that altered this ratio through manipulations in ploidy, injection of exogenous DNA, removal of cytoplasm, or changes in cell size (Fig. 1A) (1, 11–18).

**Figure 1.**
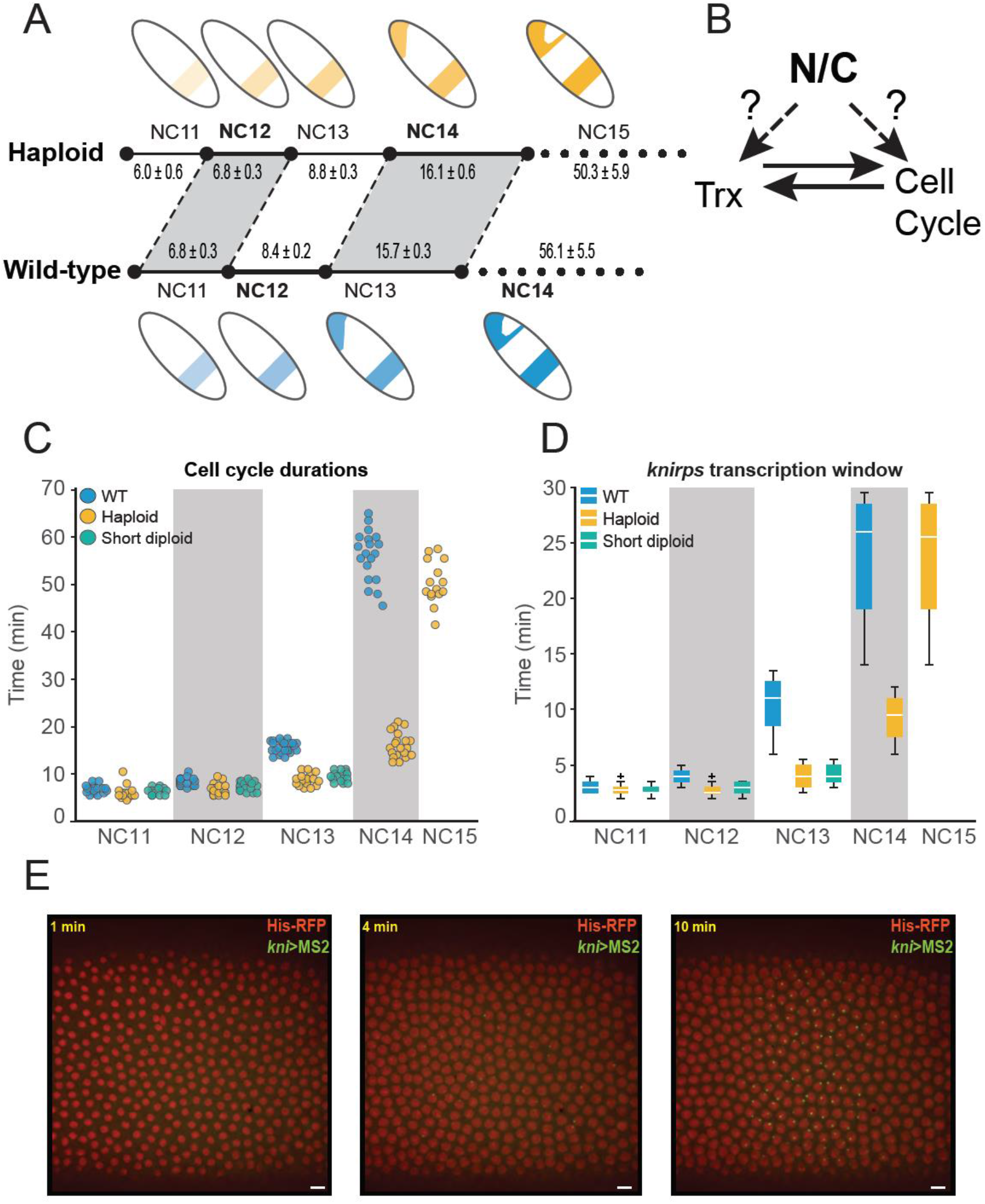
The nuclear to cytoplasmic (N/C) ratio regulates cell cycle and transcription duration. (A) Cell cycle elongation and transcription activation (as illustrated by a cartoon of the *knirps* expression pattern) are both delayed by reduction of the N/C ratio. Haploid embryos undergo one additional fast cell cycle to restore the correct N/C ratio before slowing, and all previous nuclear cycles (NCs) are correspondingly shortened. Transcription is similarly delayed. The mean cell cycle durations ± the SEM are given (in minutes) for both genotypes. (B) Since the N/C ratio affects both cell cycle and transcription it is difficult to disentangle which event is upstream, or if both sense the N/C ratio independently. (C) Scatterplot of cell cycle duration from WT (blue), haploid (yellow), and short-cycle diploid (green) embryos illustrate how ploidy affects the length of interphase and therefore the maximum potential transcriptional window. The number of embryos analyzed in WT, haploid, and short-cycle diploid in each NC is as follows: NC11 [23, 17 16]. NC12 [24,20,16], NC13 [24,20, 16]. NC14 [19, 20, N/A]. NC15 [N/A, 15, N/A]. (D) Boxplots showing that *kni* transcription duration is longer in WT than haploid, and short-cycle diploid embryos. Boxplots show minimum (10%), lower (25%), median, upper (75%), and maximum (90%) quantiles. Outliers are shown as ‘+’. The number of nuclei analyzed in (D) is as follows: 47 NC11, 184 NC12, 658 NC13, and 1051 NC14 nuclei from 6 replicate *kni>MS2* WT embryos. 42 NC11, 121 NC12, 393 NC13, 755 NC14, and 1218 NC15 nuclei from 4 replicate *kni>MS2* haploid embryos. 25 NC11, 68 NC12, and 267 NC13 nuclei from 4 replicate *kni>MS2* short-cycle diploid embryos. (E) Example images of MS2-foci in an embryo expressing *kni*>*MS2* reporter. Nuclei are marked with His2Av-mRFP. Images show *kni>MS2* expression at 0, 4, and 10 minutes after the onset of NC14. Histogram was adjusted for visualization purposes only, and images were rotated to orient the embryo (left-anterior, right-posterior).

Since transcription and the cell cycle progression are tightly coupled, disentangling which is upstream during the MBT has remained difficult (6, 19). On the one hand, transcript accumulation is necessarily dependent on the length of the transcriptional window, i.e. interphase duration (5, 20, 21). Indeed, artificial manipulation of the cell cycle results in corresponding changes to the timing of ZGA in *Xenopus*, zebrafish, and *Drosophila* (22–25), while pharmacological inhibition of transcription does not affect cell cycle behavior in *Xenopus* or zebrafish (26–29). In *Drosophila* and zebrafish, it has been observed that the early interphases simply are not long enough to sustain robust transcription of most genes (20, 21, 30–33). On the other hand, ZGA has been implicated as upstream of cell cycle slowing. Premature activation of transcription does lead to earlier cell cycle slowing in *Drosophila*, while transcription inhibition leads to delayed cycle lengthening (in contrast to *Xenopus* and zebrafish) (34–36).

Given this complex interdependence between the cell cycle and transcription, the question of whether the N/C ratio directly or indirectly affects transcription has remained unanswered (Fig. 1B). *In vitro,* at least one transcript is directly sensitive to the N/C ratio in cell cycle arrested *Xenopus* egg extracts, but it is unclear if this direct relationship is maintained *in vivo* for any or all genes (37). *In vivo*, manipulations in ploidy coupled with RNA-seq, microarrays, or qPCR have found that haploid *Drosophila*, *Xenopus*, and zebrafish embryos have reduced gene expression when compared to their wild type counterparts with a spectrum of N/C-dependence across transcribed genes (15, 18, 38). Yet, such sampling based experiments are ill suited to determine if the observed changes in transcription are a direct response to the altered N/C ratio or an indirect response to changes in cell cycle duration, because they lack the temporal resolution required to properly account for the cumulative changes in interphase length. Moreover, examination of endogenous genes in ploidy manipulated embryos are inherently confounded by the inevitable effect on template availability. Recent, carefully designed experiments have attempted to account for this effect by making the assumption that halving the template should result in half the output (18). However, this assumes that the template is the sole limiting factor for transcription, which may not be the case for all genes (39–42). In addition, whole embryo sequencing based approaches also destroy the spatial information within an embryo, which is important since many of the early genes are spatially patterned. Fixed tissue imaging-based approaches such as *in situ* hybridization or labeled ribonucleotide incorporation can circumvent the latter issue, but have limited temporal resolution to fully address if and how transcriptional dynamics respond to the N/C ratio.

In this study, we have employed the MS2-MCP system to directly and quantitatively investigate the effects of the N/C ratio on real-time transcriptional dynamics in the early *Drosophila* embryos, in single cells. This system allows us to follow the transcription of candidate genes over the course of several cell cycles with <30 second temporal resolution. We find that for all of the genes studied, N/C-ratio-dependent changes in interphase duration result in proportional changes in total transcription output within a given cycle. For some genes such as *knirps (kni)* and *snail (sna)*, the change in cell cycle duration is the only mechanism by which the N/C ratio impinges on transcriptional output. However, other genes directly sense the N/C ratio through one of two distinct mechanisms independent of the interphase length. Some genes, such as *giant (gt)* and *bottleneck (bnk)* alter the kinetics of transcription activation such that transcription is slower at lower N/C ratios. On the other hand, the cell cycle regulator, *frs* reduces the probability of transcription initiation such that fewer nuclei are activated in haploid embryos of the same cycle. We conclude that while all genes are affected by cell cycle duration, some genes including *gt*, *bnk,* and *frs* contain regulatory elements that respond directly to the N/C ratio. Moreover, different N/C ratio responsive genes employ different mechanisms to regulate transcription in response to ploidy.

## Results

### Cell cycle duration controls the transcriptional window which determines total mRNA output

To investigate how the N/C ratio regulates transcription, we visualized nascent mRNA production in developing embryos through the MS2-MCP-based live imaging system (43–47). When the inserted 24 MS2 repeats are transcribed, the nascent RNA is bound by maternally provided MCP:GFP, allowing us to detect nascent transcripts (Fig. 1E) and the integral of fluorescent intensity over time is used as a proxy for cumulative mRNA production (47, 48). To remove the confounding effects of copy number on transcriptional output, we compared wild-type (WT) embryos that were heterozygous for a given MS2 construct to haploid embryos which also contained only a single copy of the construct (see SI Appendix for details).

Since *Drosophila* forgo cytokinesis until the MBT when the resulting syncytium is cellularized, the first 13 divisions are referred to as nuclear cycles (NCs, not to be confused with the N/C ratio). We used the *sesame/Hira185b* (*ssm185b*) mutation to generate haploid embryos (referred to hereafter as haploids) (49–51). The cell cycle lengths of haploids are shifted by one nuclear cycle (e.g., duration of diploid NC13 ≈ haploid NC14) and undergo one additional nuclear division in order to reach the same N/C ratio (where N is proportional to the total amount of DNA, not number of nuclei) as their diploid counterparts before the MBT (Fig. 1A and 1C) (11, 15, 52–55). We found that the duration of active transcription scales directly with cell cycle length for all genes studied (*kni, sna, gt, bnk,* and *frs*) (Fig. 1C, 1D and SI Appendix - Fig. S1). Regardless of each gene’s previous N/C ratio categorization (15), the change in the duration of the cell cycle and hence, the transcriptional window, had a profound effect on the total transcriptional output (SI Appendix – Fig. S2). For example, the cumulative *kni* mRNA output of each haploid cycle is delayed by precisely one cycle as compared to WT embryos (Fig. 2A and SI Appendix - Fig. S2A).

**Figure 2.**
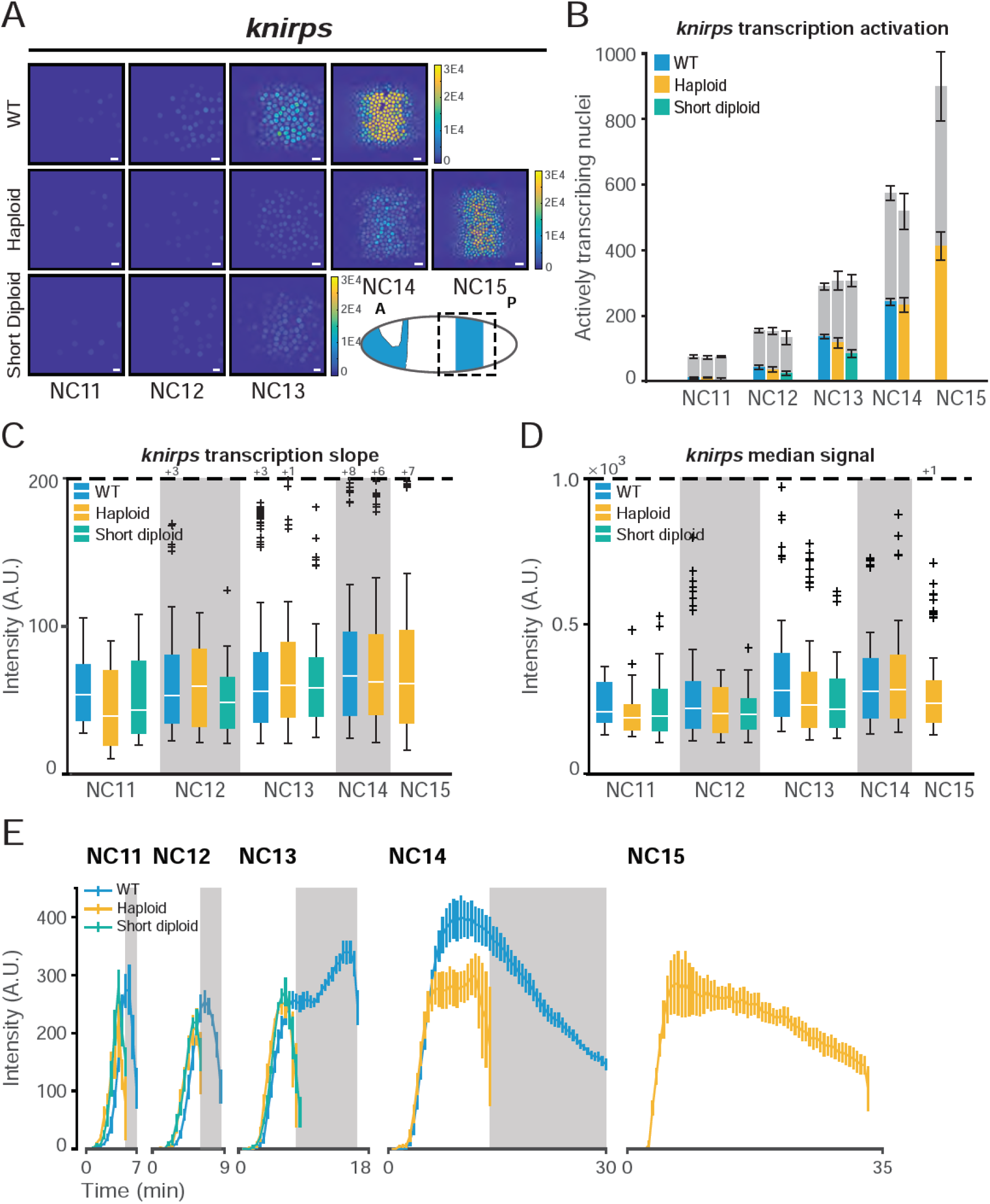
N/C ratio regulates *knirps* transcription by shortening the cell cycle. (A) Heat map showing that total RNA output is greater in representative WT embryos expressing *kni>MS2*, in NC11-NC14, than haploid or short-cycle diploid. Haploids catch up as cell cycle slows in NC15. Color bar represents total cumulative output per nucleus per NC (A.U.). The cartoon shows endogenous *kni* pattern at NC14 and dashed box indicates the area under analysis. Scale bar represents 10 µm. (B) Bar chart showing that the number of nuclei transcribing *kni>MS2* in a given nuclear cycle is similar between WT (blue), haploid (yellow), and short-cycle diploid (green) embryos. Gray bars represent the total number of nuclei analyzed in each cycle and colored bars represent the number of active nuclei. Values are similar across all genotypes. Data represented as mean ± SEM of 6 replicate *kni>MS2* WT, 4 replicate *kni>MS2* haploid, and 4 replicate *kni>MS2* short-cycle diploid embryos. (C) Boxplots showing no difference in the rate of transcriptional activation of *kni>MS2* for all actively transcribing nuclei for WT, haploid, and short-cycle diploid embryos. The rate was quantified by measuring the slope of transcription initiation. Dashed line represents cut-off for outlier values. Number of outlier values above the cut-off are given after ‘+’. (D) Boxplots showing median transcriptional activity of *kni>MS2* from all transcribing nuclei. There is a small decrease in the median for both haploid and short-cycle diploid indicating that this is an indirect effect of cell cycle duration rather than a direct effect of ploidy. (E) Average *kni*>MS2 transcriptional trajectory over time for all transcribing nuclei per nuclear cycle. Data represented as mean ± SEM. Gray boxes represent haploid mitoses. The number of nuclei analyzed in each NC and for each genotype in panel (B-E) is the same as shown in Figure 1D.

The shorter cell cycle in haploids makes it difficult to disentangle the effect of cell cycle duration versus N/C ratio on transcription. To quantify changes in transcriptional activity independent of cell cycle length, we compared haploids to diploid embryos with similarly shortened cell cycles produced by mutation of checkpoint kinase 1 (*grp/chk1*, referred to hereafter as short-cycle diploids) (56–58). These embryos undergo the early nuclear cycles normally, but fail to slow the cell cycles leading up to the MBT, resulting in a catastrophic 13^th^ mitosis, and therefore cannot be analyzed after NC13 (58). We found that interphase length was similar between haploids and short-cycle diploids, allowing us to compare the direct effect of N/C ratio on transcription (Fig. 1D). The total mRNA output for *kni* and *sna* was comparable between haploids and short-cycle diploids, suggesting that the transcription of these genes is primarily affected by cell cycle length (Fig. 2 and SI Appendix – Fig. S2A and S2B). However, not all genes responded to haploids and short-cycle diploids in an N/C-ratio-independent manner (see below).

We next asked if the observed N/C ratio effects on transcription output could be attributed solely to the change in cell cycle length or if any of these effects were direct results of N/C ratio sensing. If the regulatory elements of a given gene directly sense the N/C ratio, they could change the rate of polymerase loading, the rate of elongation, and the median transcriptional signal, irrespective of cell cycle length. Or, the timing and/or probability of gene activation could be affected based on the respective N/C ratio. To this end, we measured three parameters of transcriptional activity that are independent of the cell cycle duration: the number of actively transcribing nuclei, the rate of transcriptional activation (slope), and the median amplitude of transcription in each cell cycle. *kni* and *sna* behaved in a purely time dependent manner with no evidence of direct N/C ratio effects on transcription. The number of active nuclei and the rate of transcription activation within a given cell cycle were indistinguishable between WT, haploid, and short-cycle diploid (Fig. 2B and 2C, SI Appendix – Fig. S3). There was a small decrease in the median amplitude of transcription in haploid cell cycles (Fig. 2D).

However, since the signal in the early cycles never reaches steady state, we reasoned that the shortened cell cycle in haploids may prematurely truncate transcription during the rising phase. Indeed, the average fluorescent signal across all transcribing nuclei over time confirmed a premature termination of transcription in both haploids and short-cycle diploids (Fig. 2D and 2E). When fluorescent signal was aligned by equivalent N/C ratio, and therefore equivalent cell cycle length between WT and haploids (e.g. WT NC13 and haploid NC14), the transcriptional trajectories were better aligned, suggesting that cell cycle duration plays a major role in transcriptional regulation of *kni* and *sna* (Fig. 2E and SI Appendix – Fig. S3E, S4A and S4B).

We also noted that all genotypes displayed a small increase in the median signal with age, consistent with transcriptional memory or increased translation of transcriptional activators (42, 59, 60). This indicates that the small change in the transcriptional amplitude observed between WT and haploids can be attributed to changes in cell cycle duration or cell cycle state, not changes in the underlying competency of the transcription machinery in response to the N/C ratio. Therefore, for one category of genes, represented by *kni* and *sna*, the only perceivable effect of the N/C ratio on transcriptional output is due to changes in cell cycle length.

### The N/C ratio controls transcriptional kinetics for a subset of genes

In contrast to *kni* and *sna*, the effects of the N/C ratio on other genes cannot be explained solely by changes in cell cycle durations. We found two categories of direct N/C-dependent gene behavior. In the first, the N/C ratio regulates the kinetics of transcription activation. The gap gene *gt* and the cellularization gene *bnk* are examples of such regulation. Similar to *kni* and *sna*, *gt* and *bnk* displayed a delayed accumulation of nascent RNA with each nuclear cycle (Fig. 3A, SI Appendix – Fig. S2C, S2D, and S5). Additionally, there was no delay in global activation of these genes, as WT and haploids both showed comparable numbers of actively transcribing nuclei in each cell cycle (Fig. 3B, SI Appendix – Fig. S5B). However, unlike *kni* and *sna*, where cell cycle duration was the main factor of N/C-ratio-dependent changes in transcription, *gt* and *bnk* expression in haploids demonstrated a decreased rate of transcriptional activation compared to both WT and short-cycle diploids in the early cell cycles (Fig. 3C and SI Appendix – Fig. S5C). This N/C-ratio-dependent initial trajectory suggests that the kinetics of *gt* and *bnk* transcription initiation are directly responsive to embryo ploidy. The median signal further confirmed that the initial rate of transcription and average transcriptional amplitude were lower in haploids in the early cell cycles (Fig. 3D and SI Appendix - Fig. S5D). The reduction in median transcriptional amplitude was also observed in the average nuclear trajectories, although the nuclei-to-nuclei variability in the timing of transcription activation obscured this distinct difference (Fig. 3E and SI Appendix – S5E). This result suggests that the kinetics of polymerase loading are directly sensitive to the N/C ratio for a class of genes. Surprisingly, we observed a much higher *gt* signal in the short-cycle diploids compared to haploids and WT (Fig. 3D and 3E). This may be due to the faster replication, and therefore greater early template availability in the short cell cycle conditions, or some other, more direct effect of *chk1* activity, though this effect is not as striking in *kni*, *bnk*, or *frs*. Nonetheless, the fact that the short-cycle diploid is more, not less, active than WT strongly indicates that the reduced rate of transcriptional activation observed in haploids is not a result of cell cycle duration alone.

**Figure 3:**
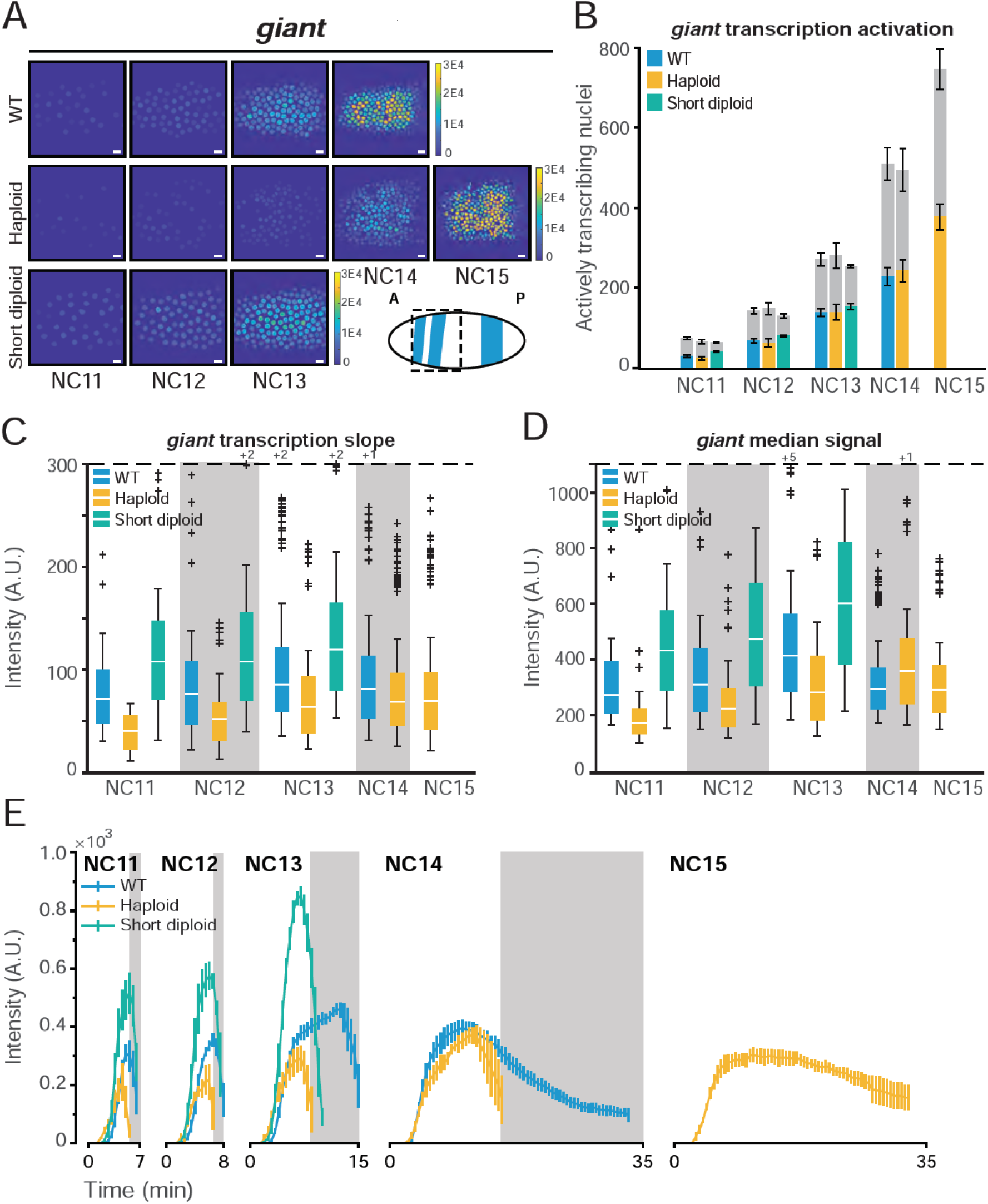
*gt* responds to the N/C ratio in a kinetic-dependent manner. (A) Heat map of total RNA output for a representative embryo expressing *gt>MS2*. RNA output is comparable between WT and short-cycle diploid, and is lower in haploids. Color bar represents total cumulative output per nucleus per NC (A.U.). The cartoon shows endogenous *gt* pattern at NC14 and dashed box indicates the area under analysis. Scale bar represents 10 µm. (B) Bar chart showing that the number of nuclei transcribing *gt>MS2* in a given nuclear cycle is similar between WT (blue), haploid (yellow), and short-cycle diploid (green) embryos. Gray bars represent the total number of nuclei analyzed in each cycle and colored bars represent the number of active nuclei. Data represented as mean ± SEM of 6 replicate *gt>MS2* WT embryos, 5 replicate *gt>MS2* haploid embryos, and 4 replicate *gt>MS2* short-cycle diploid embryos. (C) Boxplots showing the rate of transcriptional activation (initial slope) activation of *gt>MS2* for all actively transcribing nuclei for WT, haploid, and short-cycle diploid embryos. The initial slope is lower in haploids compared to WT. (D) Boxplots showing median transcriptional activity of *gt*>*MS2* from all transcribing nuclei. The average amplitude of transcription is reduced in haploids. (E) Average *gt*>MS2 transcriptional trajectory over time for all transcribing nuclei per nuclear cycle. Data represented as mean ± SEM. Gray boxes represent haploid mitoses. The number of nuclei analyzed in (B-E) is as follows: 164 NC11, 362 NC12, 730 NC13, and 1077 NC14 nuclei from 6 replicate *gt>MS2* WT embryos. 80 NC11, 270 NC12, 587 NC13, 1013 NC14, and 1513 NC15 nuclei from 5 replicate *gt>MS2* haploid embryos. 158 NC11, 308 NC12, and 581 NC13 nuclei from 4 replicate *gt>MS2* short-cycle diploid embryos.

### The N/C ratio regulates binary activation probability for a subset of genes

The second category of direct N/C ratio responsive transcription behavior was more striking than the change in the rate of transcription observed in *gt* and *bnk*. In this class, which consists of the cell cycle regulator *frs*, the global probability of gene activation was directly N/C ratio sensitive. While *kni, sna, gt,* and *bnk* showed gradual activation from NC11 to NC14, *frs* showed a dramatic switch in the number of active nuclei between cell cycles in WT, going from 16% in WT NC12 to 80% in WT NC13 (Fig. 4A and 4B). This is consistent with the sharp increase in RNA observed during NC13 by time-course RNA-seq (61, 62). We found the switch from majority inactive to majority active nuclei was delayed by one cell cycle in haploids, occurring between NC13 and NC14 rather than between NC12 and NC13 (Fig. 4A and 4B). Notably, the number of active nuclei were comparable between WT and in short-cycle diploids in NC13 (Fig. 4B), demonstrating that the switch-like activation of *frs* is a direct effect of ploidy, as opposed to cell cycle duration. During haploid NC14, *frs* was fully activated and the fraction of actively transcribing nuclei stayed constant in haploid NC15, as we observed in the other genes (Fig. 4D). It is important to note that ploidy does not change the total number of nuclei in a given cycle, and only alters the amount of DNA in each nucleus (Fig. 3B and 4B).

**Figure 4.**
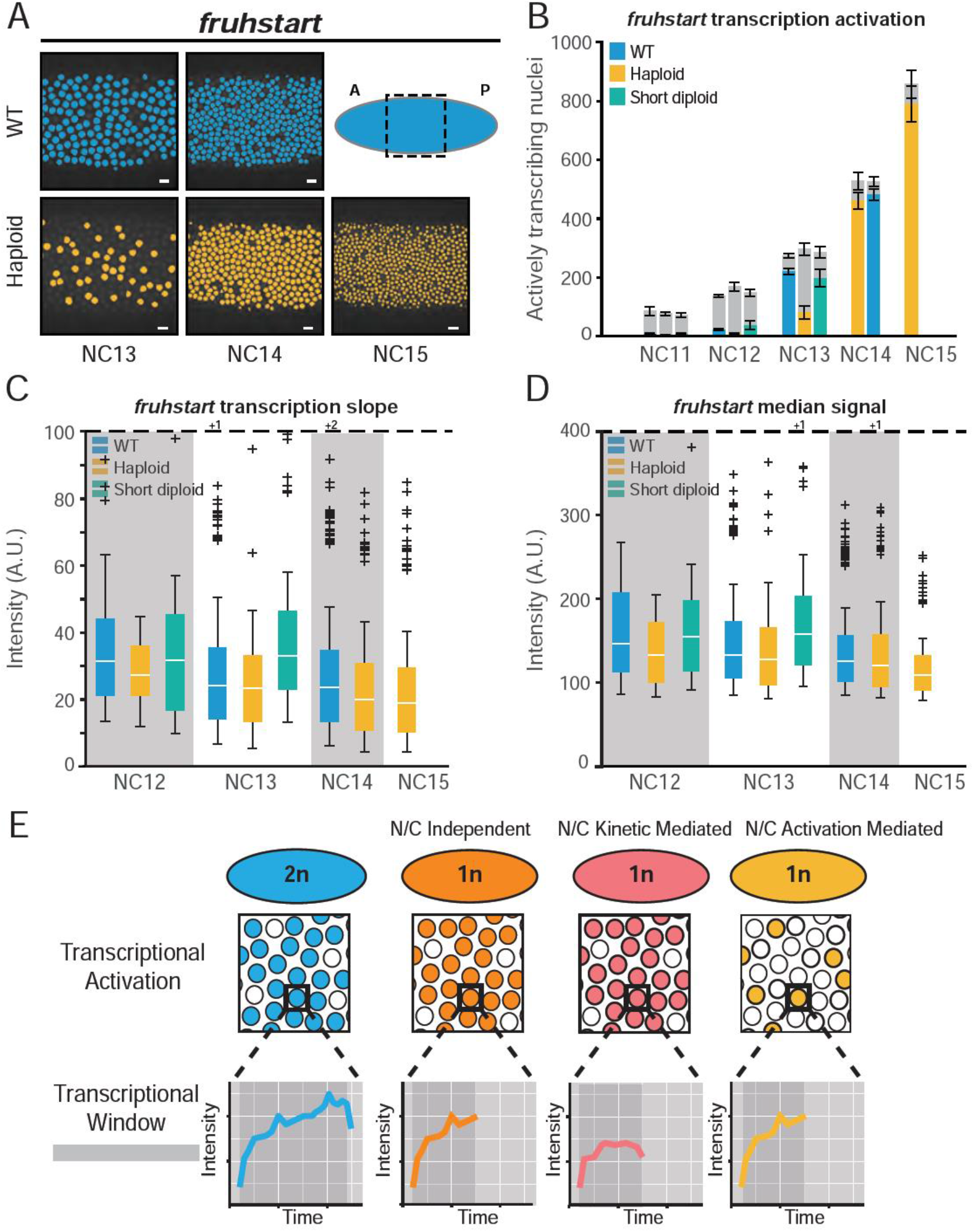
The N/C ratio regulates *frs* in an activation-mediated manner. (A) *frs* nuclei that ever showed an active transcription are false colored in blue (WT) from NC13-NC14 and yellow (haploids) from NC13-NC15. Note the decreased number of active nuclei in haploid NC13. The cartoon shows endogenous ubiquitous *frs* pattern at NC14 and dashed box indicates the area under analysis. Scale bar represents 10 µm. (B) Bar chart showing a delay in the switch from mostly inactive to mostly actively transcribing *frs-MS2* in haploid (yellow) compared to WT (blue) embryos. Short-cycle diploids (green) activate similarly to WT in NC13. Gray bars represent the total number of nuclei analyzed in each cycle and colored bars represent the number of active nuclei. Data represented as mean ± SEM of 5 replicate *frs-MS2* WT, 4 replicate *frs-MS2* haploid, and 4 replicate *frs-MS2* short-cycle diploid embryos. (C) Boxplots showing that the rate of transcriptional activation (initial slope) of *frs-MS2* is similar for all actively transcribing nuclei for WT, haploid, and short-cycle diploid embryos. (D) Boxplots showing median transcriptional activity of *frs-MS2* from all transcribing nuclei. The average transcription amplitude is comparable among WT, haploid, and short-cycle diploid embryos. The number of nuclei analyzed in (B-D) is as follows: 55 NC12, 791 NC13, and 1921 NC14 nuclei from 5 replicate *frs-MS2* WT embryos. 17 NC12, 177 NC13, 1355 NC14, and 1510 NC15 nuclei from 4 replicate *frs-MS2* haploid embryos. 97 NC12, and 546 NC13 nuclei from 4 replicate *frs-MS2* short-cycle diploid embryos. (E) Model depicting the different modes by which N/C ratio modulates transcription. The WT (2n) for all the genes has a number of actively transcribing nuclei each with a trace over a period of time, indicated by the generic curve. In the N/C independent genes, the number of active nuclei and the rate of transcription remains identical to WT; however due to a shortened cell cycle duration, transcription is terminated earlier than WT. In N/C kinetic-mediated genes, while the number of nuclei activated are similar to WT, the rate at which transcription occurs is slower than WT. In N/C activation-mediated genes, the number of nuclei activated is significantly lower than WT, while the rate of transcription remains comparable to WT.

For those nuclei that are active in NC13 haploids and short-cycle diploids, the total per nucleus output of *frs* is highly reflective of cell cycle duration, similar to other genes in this study (SI Appendix - Fig. S2E). This suggests that the N/C ratio may regulate only the probability of activation, rather than affecting the entire transcriptional kinetics. Indeed, the rate of transcriptional activation and the median amplitude of transcriptional activity in those nuclei that do initiate transcription in the earlier cycles are unaffected by ploidy (Fig. 4C and 4D). These results indicate that the regulatory elements of *frs* are directly responsive to the N/C ratio in a binary on/off manner and that the N/C ratio has no additional detectable effects on transcription kinetics.

## Discussion

Here, we have shown that the total transcriptional output during the MBT is a function of the N/C ratio for all measured genes in *Drosophila* and classified three major categories of N/C ratio control. For some genes (*kni* and *sna*), the primary, if not only, effect of the N/C ratio is due to N/C-dependent changes in the length of the cell cycle and hence, the length of the transcriptional window. This results in a reduction in total mRNA production within a given cycle due to the abortion of transcription at mitosis (20, 21, 33, 63). However, the activation of other genes (*gt, bnk,* and *frs*) is directly sensitive to the N/C ratio, regardless of the cell cycle duration. We show that this direct sensing of N/C ratio is manifested as two different gene expression regulatory paradigms: 1) altering the kinetics of transcriptional activation of actively transcribing nuclei, and 2) regulating the probability of gene activation in a given cell cycle (Fig. 4E).

The existence of the first category of N/C ratio sensing, which affects solely the rate of transcription without changing the number of actively transcribing nuclei per cycle, suggests that the *cis* regulatory elements of these genes directly respond to the N/C ratio. In particular, the rate of polymerase elongation during transcription or the rate of polymerase recruitment to the promoter is likely affected in haploids, independent of cell cycle length. We found that such N/C kinetic-mediated genes like *gt* and *bnk* showed faster rates of transcription in WT compared to other genes, implying that the rapidly transcribing genes may be more sensitive to the kinetic-mediated transcription regulated by the N/C ratio (SI Appendix – Fig. S6).

In contrast, the activation probability N/C ratio sensing gene, *frs*, is uniquely switch-like in our study. We speculate that this switch-like behavior may be a consequence of its responsiveness to the exponentially increasing N/C ratio (Fig. 4A and 4B). This process may set a threshold for the recruitment of polymerase to the *frs* promoter, or, more likely, its release from pausing at the *frs* promoter. The promoter of *frs,* as well as other genes we assayed, are already nucleosome-free by NC11 indicating chromatin opening alone cannot explain their differences in timing (54). Nonetheless, *frs* is responsive to experimental alterations in histone availability which is decreasing over the course of the early divisions (55, 62). This is in contrast to the other zygotic genes that do not respond directly to the N/C ratio (*sna* and *kni*), and also have limited histone sensitivity (62). Therefore, if histone abundance is regulating zygotic genome activation, it must do so through a gene specific mechanism and not via a global change in accessibility.

We also note that most manipulations of the N/C ratio are, in fact, manipulations in ploidy. In these cases, since the amount of template for a given transcript is altered, we speculate that the transcript accumulation is much more dramatically affected than a simple response to the change in N/C-ratio-dependent cell cycle duration. Template reduction could also directly affect the process of cell cycle slowing. For example, in the model where RNA-polymerase on the DNA acts as a source of replication stress, simply halving the amount of DNA without changing other aspects of transcription would halve the number of such conflicts embryo-wide (58). This may permit an additional cell cycle in haploids to allow for the critical number of global conflicts to induce the checkpoint response to slow the cell cycle. In sum, our findings constrain the available models that attempt to decipher how the N/C ratio influences both the cell cycle and transcription, directly or indirectly.

## Data Availability

Detailed materials and methods, and all other data discussed in the manuscript are available in the SI Appendix.

## Acknowledgments

We are grateful to Gary Laevsky and the Molecular Biology Confocal Imaging Facility and Gordon Gray and the *Drosophila* Media Core Facility at Princeton University for technical support. We thank Michal Levo for reagents. We thank Eric Wieschaus, Stas Shvartsman, and Mike Levine for discussion. Stocks obtained from the Bloomington *Drosophila* Stock Center (NIH P40OD018537) were used in this study. SS and BL are supported by NIH R35GM133425.

## Supplementary Information Text

### Materials and Methods

#### Generation of wild-type, diploid embryos

Wild-type embryos were produced by crossing *y*,*w*;;MCP:GFP,His2Av-mRFP virgin females to *y*,*w*;; males. The resulting embryos were provided with one maternal copy of the MCP:GFP,His2Av-mRFP while the desired MS2 was provided paternally. Crosses were conducted in collection cups at 25°C. We observed no difference in MS2 signal between wild-type embryos that had MS2 provided maternally and from those provided paternally.

#### Generation of haploid embryos

Haploid embryos were derived from the *sesame/Hira^185b^* (*ssm*) line. Haploid embryos were made by crossing hemizygous *ssm* mutant males in the desired MS2 background to heterozygous *ssm*/FM7c virgin females in the MCP:GFP,His2Av-mRFP background to produce *ssm* homozygous females that were heterozygous for both MCP:GFP and the desired MS2. These females, in turn lay embryos that do not include the male contribution and are thus haploid. For example, to observe haploid *sna* transcription: *w,ssm*;;*sna-*Promoter-Distal-Enhancer-MS2males were crossed to *w,ssm*/FM7c;;MCP:GFP,His2Av-mRFP virgins. Homozygous *ssm* females were collected, crossed with *y*,*w*;; males and placed in collection cups at 25°C. Embryos laid by the *ssm* homozygous mothers with one copy of the MCP:GFP,His2Av-mRFP and one copy of the desired MS2 were used for haploid. Only embryos that inherited the desired MS2 were analyzed.

#### Generation of short-cycle diploid (*grp*) embryos

Short-cycle diploid embryos were derived from *w*; *grp^1^* / CyO; *ry^506^.* Short-cycle diploid embryos were produced by crossing *w*; *grp^1^* / CyO; *ry^506^* virgins to *w*; *grp^1^* / CyO; MCP:GFP,His2Av-mRFP males. Resulting *grp^1^* homozygous virgins were collected, crossed to desired MS2 males, and placed in collection cups at 25°C. Embryos laid by these mothers were used for short-cycle diploid embryos.

##### MS2 constructs

###### *kni*-vk33

The *knirps*>*MS2-yellow* reporter was constructed by amplifying the *knirps* regulatory region from 5.5 kb upstream to 1kb downstream of the promoter, including the *knirps* promoter. This sequence was cloned in the pBphi-*MS2-yellow* vector (1) using the NotI and BamHI restriction sites.

###### *gt*-vk33

The *giant*>*MS2*-*yellow* reporter was generated using the same method as *knirps* to clone the *giant* regulatory region from 10kb upstream of the promoter including the *giant* promoter.

###### *bnk*-vk18

The *bottleneck>MS2-yellow* reporter was generated using the same method as *knirps* to clone the *bottleneck* regulatory region 206 bp upstream and 48 bp downstream of the *bottleneck* transcription start site (2).

###### Endogenous *frs*-MS2

*frs*-MS2 was generated using CRISPR-cas9 homology-directed repair to insert 24x MS2 loops into the 5′UTR of the endogenous *frs* locus. A single target site within the *frs* 5′UTR was selected using the Target Finder (http://targetfinder.flycrispr.neuro.brown.edu) tool (3).

gRNA oligo sequences

Sense: CTTCGCGACATAATAACTGCTAGGC

Antisense: AAACGCCTAGCAGTTATTATGTCGC

The annealed gRNA oligo was subcloned into the pU6-BbsI-chiRNA vector (a gift from Melissa Harrison & Kate O’Connor-Giles & Jill Wildonger, Addgene plasmid #45946) via BbsI restriction sites. Approximately 1-kb fragments of *frs* homology arm sequences were synthesized and inserted into the pHD-MS2-loxP-dsRed-loxP plasmid (4) (GeneWiz,Inc.). The gRNA and *frs* homology arm MS2 plasmids were co-injected into *nos*-Cas9 embryos (TH00787.N), and DsRed+ progeny were screened (BestGene). The resulting endogenous *frs*-*MS2* fly strains were homozygous viable.

##### Live imaging of transcription

All fly stocks were maintained by standard method at 25°C and were grown on standard cornmeal media. All embryos were collected on apple juice agar plates. Sex of the embryo was not considered in this study. Embryos were collected after laying for 1.5 hours at 25°C then staged as pre-blastoderm with halocarbon oil. Staged embryos were then washed with DI H2O to remove any halocarbon oil, dechorionated with 4% sodium hypochlorite for 1-1.5 minutes, mounted on a 35mm coverslip dish (MatTek), and covered with water.

All movies were acquired at 23-24°C using a Nikon Ti-E confocal microscope with a Yokogowa CSU-21 spinning disk module. Images were acquired with a Plan-Apochromat 40×1.3 NA oil objective using a 488 and 561 laser to visualize MCP:GFP and His2Av-mRFP, respectively, at a time resolution of 30s/frame. At each time point, a stack of 13 images was taken at 0.5 μm steps. The same exposure and laser settings were used for all samples. All frames were acquired as 16-bit images.

Imaging began when nuclei first emerged onto the surface of the embryo at NC10 until gastrulation (approximately 60 minutes after entry into NC14 or NC15 in haploids). Only the first 30 minutes of NC14 (wild-type) or NC15 (haploid) were analyzed due to the greater z-resolution needed to capture the total MS2 signal during cellularization. Imaging of short-cycle diploid embryos ended after the catastrophic 13th mitosis.

##### Quantification and Statistical Analysis

All the image processing methods and analyses were implemented in MATLAB (R2018b, MathWorks). Histograms of all the snapshots and movies shown in Figure 1 and in Supplemental Movies were adjusted for visualization purposes. All the analyses were performed with raw images.

##### Nuclei Segmentation and tracking

At each time frame, maximum projections for all 13 z-sections per image were obtained. His2Av-mRFP labeled channel was used to segment nuclei. Nuclei-labeled channels were first filtered with Gaussian filtering to minimize signal noise, and then were converted into binary images using a threshold value using Otsu’s method. Frames were manually corrected as needed to ensure proper segmentation. The number of nuclei that were segmented from each frame was obtained and the center of mass of each nucleus was assigned x and y coordinates for tracking. Nuclei tracking within each nuclear cycle was obtained by finding the nucleus with minimal movement across the segmented frames. In all frames, the nuclei located at the edge of the frame were excluded from the analysis.

##### MS2 signal extraction

MS2 fluorescent intensities were recorded using maximum projections of raw images and were extracted from within each nucleus after nuclei segmentation. After subtracting the background signal, the MS2 signal in a given nucleus was determined by averaging the top two pixels with the highest fluorescence intensity within each nucleus.

##### Defining active nuclei for various metrics

For all genotypes, active nuclei were defined as the nuclei that exceed an MS2 fluorescence intensity threshold of at least 50 a.u. for more than 30% of the total duration of a given cell cycle. This criterion was used to compute the various metrics and properties, including total mRNA output, duration of active transcription, and average trajectories of all cell cycles. When obtaining the number of active nuclei as a fraction of the total nuclei in a cell cycle, all nuclei that show an MS2 signal above 50 a.u. at any time point in a given cell cycle were considered.

##### Plots

Since the variability between individual nuclei within an embryo was determined to be greater than embryo to embryo variability according to T-test values, nuclei from all replicates were merged to obtain the figures. In all boxplots, the box indicates the 25% quantile and 75% quantile, and the solid line indicates the median. The top and bottom whiskers correspond to the 10th and 90th percentiles of each distribution. The error bars represent the standard error of the mean (SEM) of all replicates. Wild-type NC14 and haploid NC15 were analyzed for only 30 minutes (60 frames).

##### Trajectories of cell cycles

Transcriptional activity of each MS2-reporter gene and frs*-MS2* can be plotted over time at single-cell resolution. Each cycle was normalized to start at frame 1, or the first frame after mitosis. Segmentation and further analysis were performed 3~4 frames after mitosis to minimize nuclei movement and obtain nuclei lineages. To obtain an average transcriptional trajectory per cell cycle, data from all active nuclei of a particular genotype in a given cell cycle was combined.

##### Total mRNA output

The fluorescence intensity at a given time frame in each nucleus was used as a proxy to measure instantaneous amplitude of a given transcript. Cumulative mRNA output of a nucleus was computed by integrating an active nucleus trajectory of fluorescence intensity over time.

##### Median signal of cell cycles

The trajectories were smoothened using the local regression (LOESS) method. The maximum amplitude for each nucleus was obtained using the smooth curve. The median signal refers to the median fluorescence intensity an active nucleus exhibits during a given cell cycle.

##### Duration of cell cycles

Duration of a cell cycle was determined to be the time in between two mitosis. The duration of active transcription in a given cell cycle was the time during which the MS2 signal exceeded the predetermined threshold.

##### Transcription slopes

The transcription slope of an active nucleus was obtained by measuring the initial slope of the nucleus’s smoothened fluorescence trajectory. The smoothened curve was interpolated by a factor of 10. The transcription slope was determined to be the slope of the best-fit line after linear regression on the first 30 points above 50 a.u.

## Supplemental Figures

**Fig. S1.**
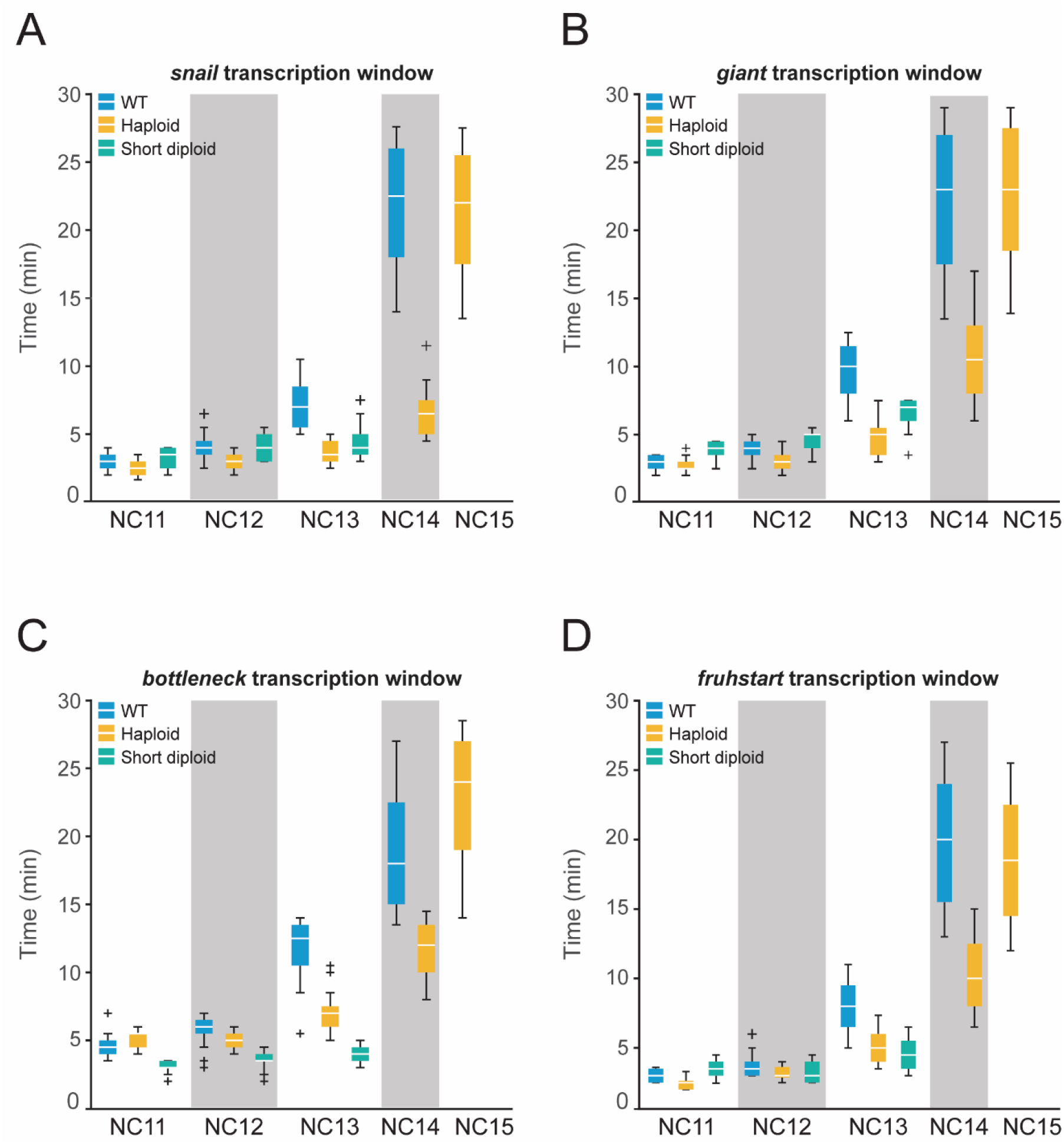
Duration of active transcription scales directly with cell cycle length. (A-D) Boxplots of transcription duration for (A) *sna>MS2,* (B) *gt>MS2,* (C) *bnk*>*MS2*, (D) *frs-MS2*, and per nucleus for WT (blue), haploid (yellow), and short-cycle diploid (green) embryos throughout the syncytial blastoderm stage. Boxplot shows minimum (10%), lower (25%), median, upper (75%), and maximum (90%) quantiles. Outliers are shown as ‘+’. The number of nuclei examined for *gt>MS2,* and *frs-MS2* is described in the legends of Figure 3E and 4D, respectively. For *sna>MS2,* the number of nuclei analyzed is as follows: 109 NC11, 163 NC12, 272 NC13, and 1283 NC14 nuclei from 4 replicate WT, 58 NC11, 153 NC12, 239 NC13, 375 NC14, and 1259 NC15 nuclei from 3 replicate haploid, and 146 NC11, 175 NC12, and 139 NC13 nuclei from 4 replicate short-cycle diploid embryos. For *bnk-MS2,* the number of nuclei analyzed is as follows: 137 NC11, 272 NC12, 497 NC13, and 726 NC14 nuclei from 3 replicate WT, 89 NC11, 333 NC12, 662 NC13, 1062 NC14, and 1196 NC15 nuclei from 4 replicate haploid, and 107 NC11, 186 NC12, and 327 NC13 nuclei from 2 replicate short-cycle diploid embryos.

**Fig S2.**
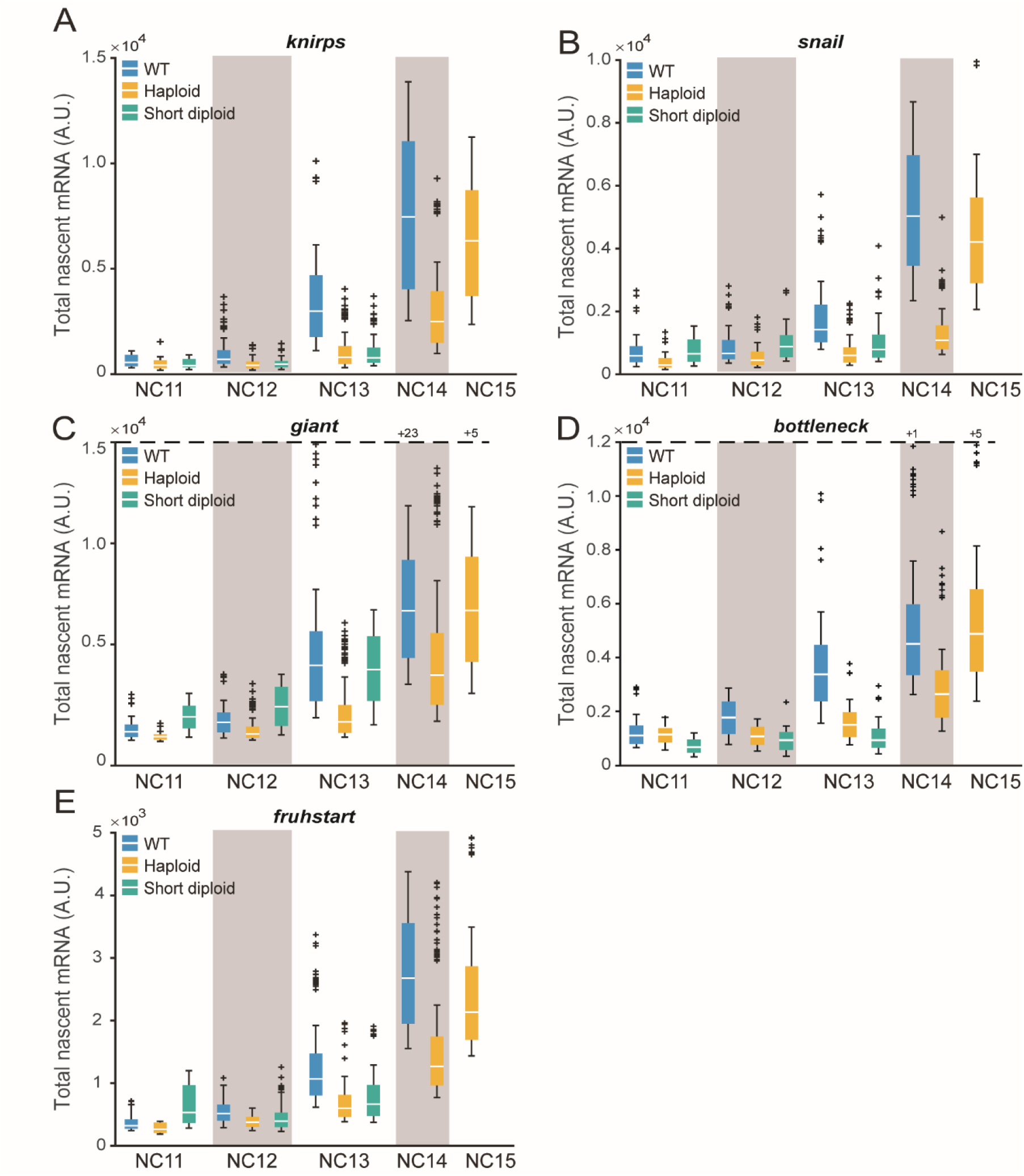
Total RNA output depends on cell cycle duration. (A-E) Boxplots of total RNA output for (A) *kni>MS2,* (B) *sna>MS2,* (C) *gt*>*MS2*, (D) *bnk-MS2*, and (E) *frs-MS2* per nucleus for WT (blue), haploid (yellow), and short-cycle diploid (green) embryos throughout the syncytial blastoderm stage. Dashed line represents cut-off for outlier values. Number of outlier values above the cut-off are given after ‘+’. The number of analyzed nuclei is the same as in Figure S1.

**Fig S3.**
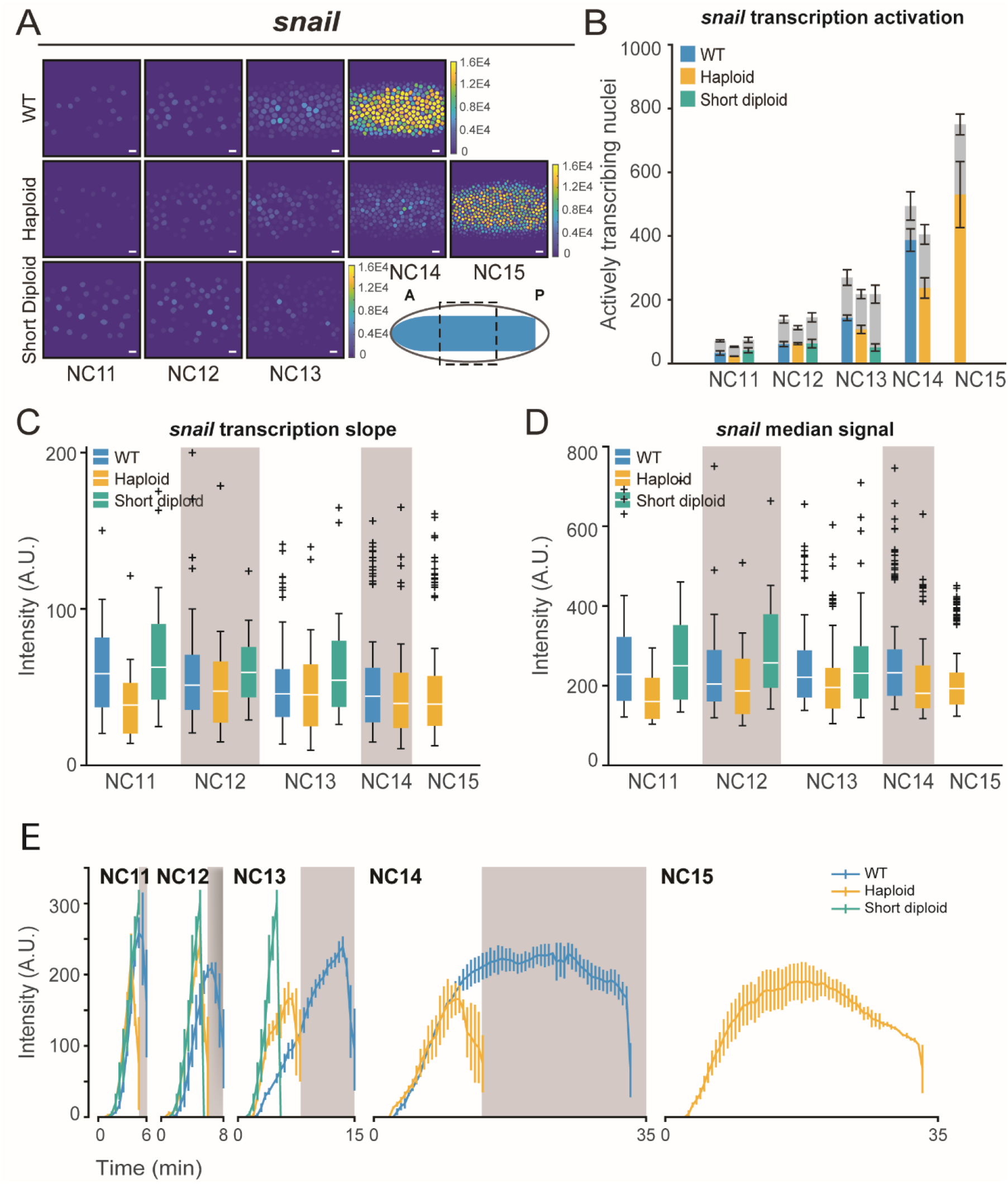
*sna* transcription is mediated mainly by cell cycle duration. (A) Heat map showing that total RNA output is greater in representative WT embryos expressing *sna>MS2*, in NC11-NC14, than haploid, or short-cycle diploid. Haploids catch up as cell cycle slows in NC15. Color bar represents total cumulative output per nucleus per NC (A.U.). The cartoon shows endogenous *sna* pattern at NC14 and dashed box indicates the area under analysis. Scale bar represents 10 µm. (B) Bar chart showing that the number of nuclei transcribing *sna>MS2* in a given nuclear cycle is similar between WT (blue), haploid (yellow), and short-cycle diploid (green) embryos. Gray bars represent the total number of nuclei analyzed in each cycle and colored bars represent the number of active nuclei. Data represented as mean ± SEM of 4 replicate *sna-MS2* WT embryos, 3 replicate *sna-MS2* haploid embryos, and 3 replicate *sna-MS2* short-cycle diploid embryos. (C) Boxplots showing comparable rates of transcriptional activation of *sna>MS2* for all actively transcribing nuclei for WT, haploid, and short-cycle diploid embryos. (D) Boxplots showing median transcriptional activity of *sna*>*MS2* from all transcribing nuclei. (E) Average *sna*>*MS2* transcriptional trajectory over time for all transcribing nuclei per nuclear cycle. Data represented as mean ± SEM. Gray boxes represent haploid mitoses. The number of analyzed nuclei in (B-E) is the same as in Figure S1 for *sna>MS2.*

**Fig S4.**
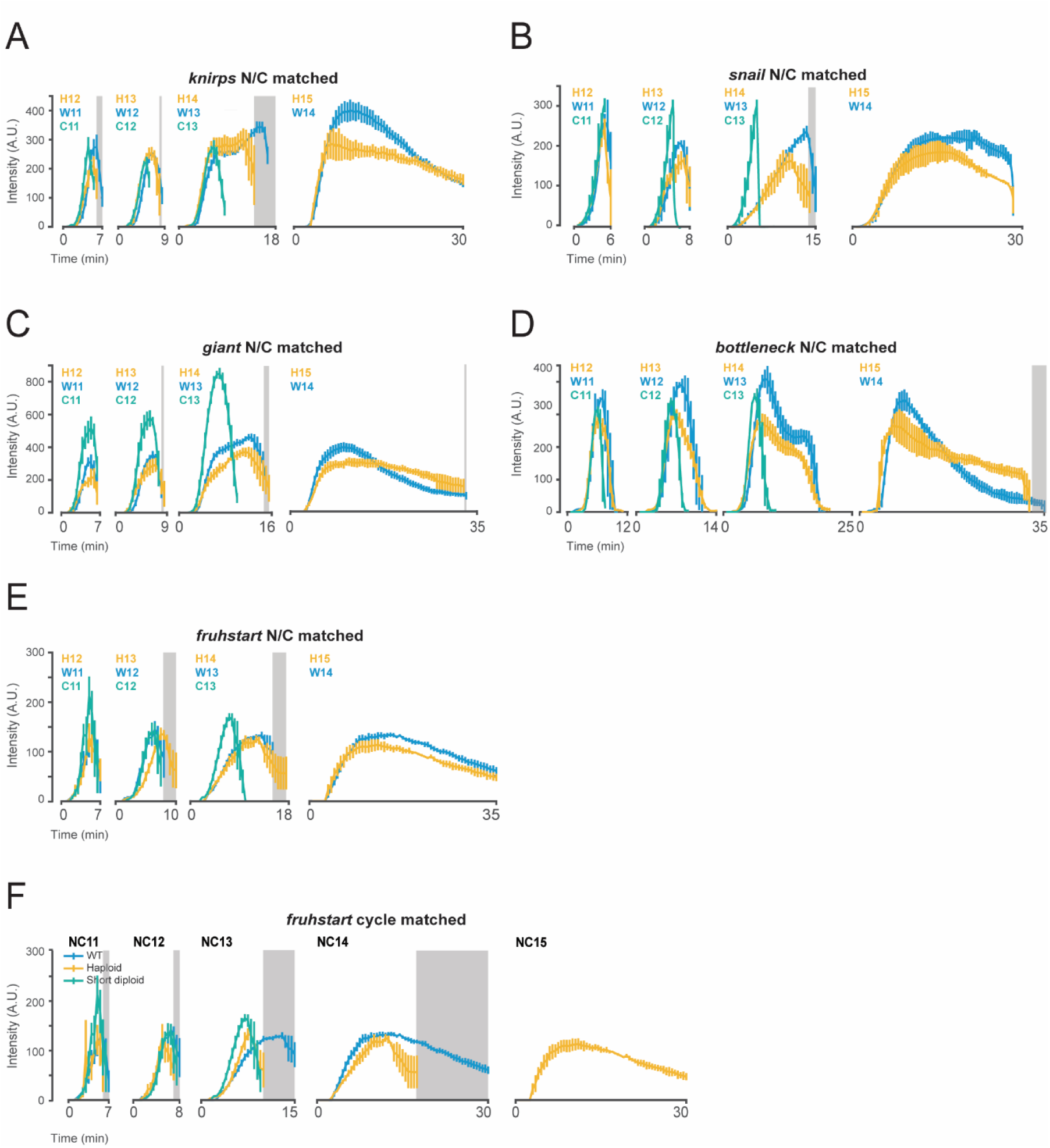
N/C ratio-matched trajectories. (A-E) Average transcriptional trajectory over time for all transcribing nuclei aligned by the N/C ratio for (A) *kni>MS2,* (B) *sna>MS2,* (C) *gt*>*MS2*, (D) *bnk-MS2*, and (E) *frs-MS2*. Data represented as mean ± SEM. Gray boxes represent mitoses. (F) Average *frs*>*MS2* transcriptional trajectory over time for all transcribing nuclei per nuclear cycle. Data represented as mean ± SEM. Gray boxes represent haploid mitoses.

**FigS5.**
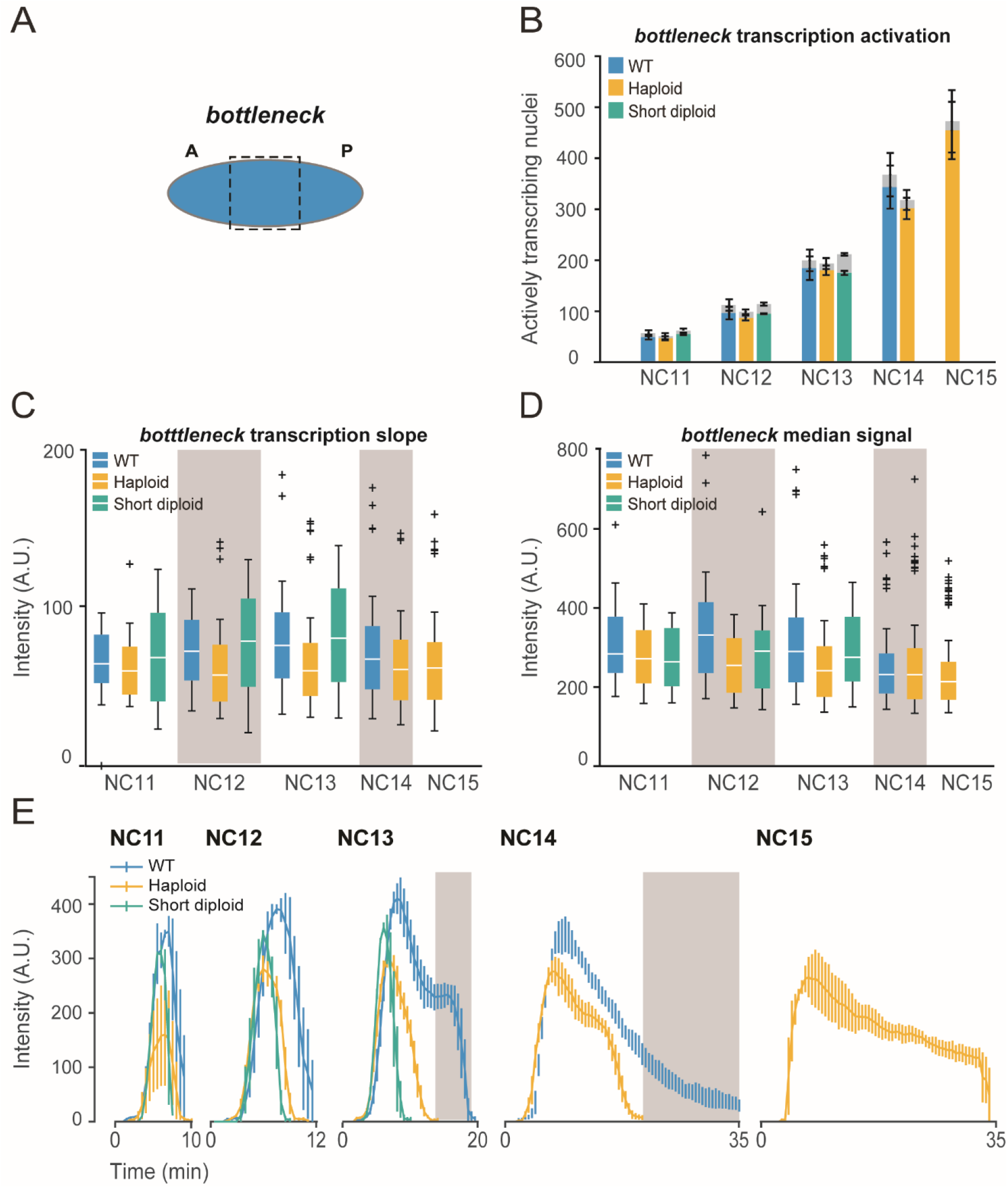
*bnk* responds to N/C ratio in a kinetic-dependent manner. (A) A cartoon that shows endogenous *bnk* pattern at NC14 and dotted box indicates the area under analysis. (B) Bar chart showing that the number of nuclei transcribing *bnk>MS2* in a given nuclear cycle is similar between WT (blue), haploid (yellow), and short-cycle diploid (green) embryos. Gray bars represent the total number of nuclei analyzed in each cycle and colored bars represent the number of active nuclei. Data represented as mean ± SEM of 3 replicate *bnk-MS2* WT embryos, 4 replicate *bnk-MS2* haploid embryos, and 2 replicate *bnk-MS2* short-cycle diploid embryos. (C) Boxplots showing the initial slope of transcriptional activation of *bnk-MS2* for all actively transcribing nuclei for WT, haploid, and short-cycle diploid embryos. The initial slope is lower in haploids compared to WT. (D) Boxplots showing median transcriptional activity of *bnk-MS2* from all transcribing nuclei. The average amplitude of transcription is reduced in haploids. (E) Average *bnk*-*MS2* transcriptional trajectory over time for all transcribing nuclei per nuclear cycle. Data represented as mean ± SEM. Gray boxes represent haploid mitoses. The number of analyzed nuclei in (B-E) is the same as in Figure S1 for *bnk-MS2.*

**Fig S6.**
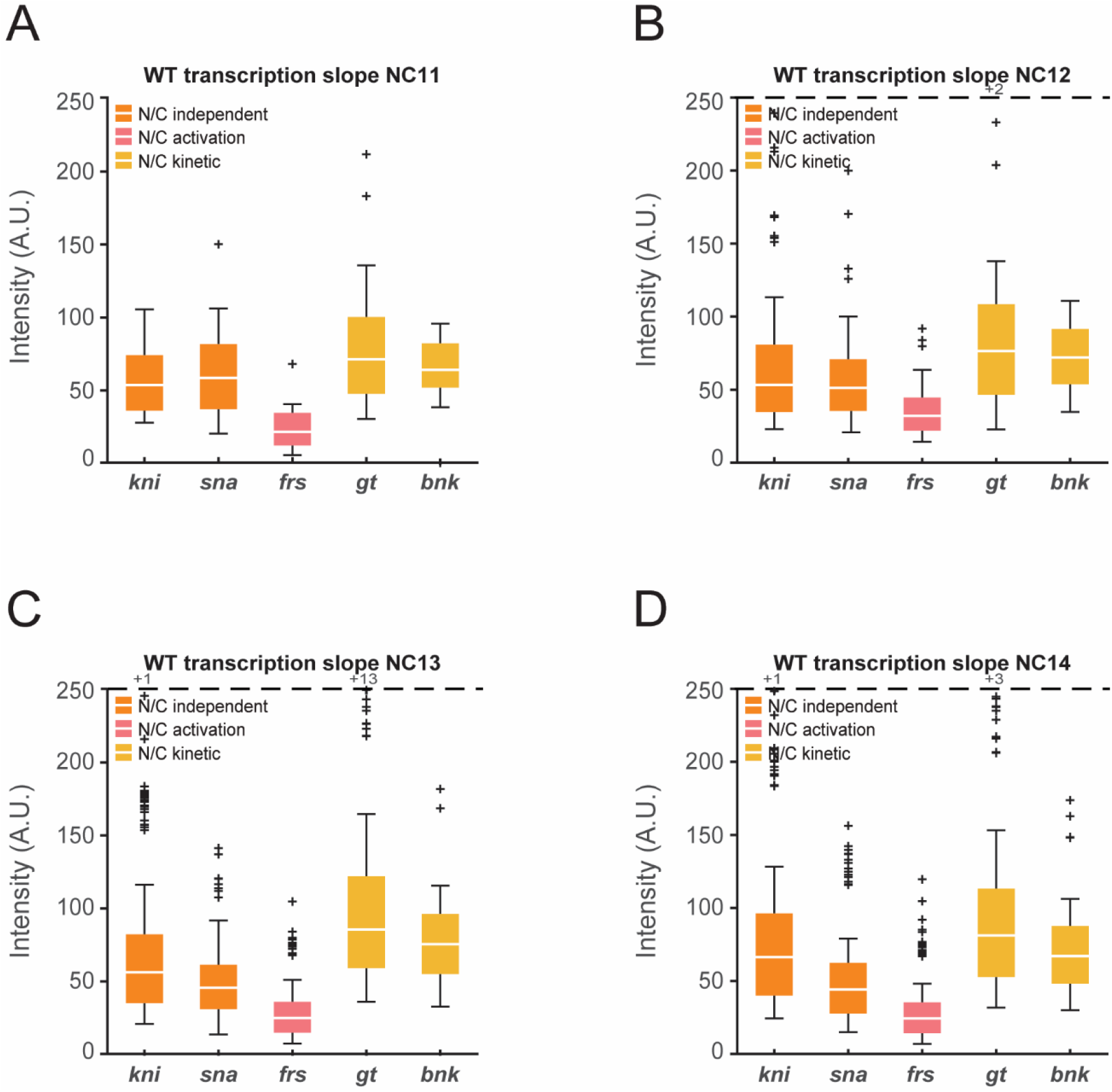
N/C-kinetic mediated genes display higher transcription slopes. (A-D) Boxplots showing the initial slopes of transcriptional activation of *kni>MS2, sna>MS2, frs-MS2, gt>MS2, and bnk-MS2* for all actively transcribing nuclei in (A) NC11, (B) NC12, (C) NC13, and (D) NC14. The genes are characterized based on the three categories: N/C independent, N/C kinetic-mediated, and N/C activation-mediated. Dashed line represents cut-off for outlier values. Number of outlier values above the cut-off are given after ‘+’.

### Supplemental Movie Legends

**Movie S1.** Live imaging of *kni>MS2*

(left) WT, NC11-NC14; (middle) haploid, NC11-NC15; (right) short-cycle diploid NC11-NC13. *MS2* signal is shown in green. Nuclei are marked with His2Av-mRFP. Histogram was adjusted for visualization purposes. Embryos are oriented top-anterior, bottom-posterior.

**Movie S2**. Live imaging of *gt>MS2*

(left) WT, NC11-NC14; (middle) haploid, NC11-NC15; (right) short-cycle diploid NC11-NC13. *MS2* signal is shown in green. Nuclei are marked with His2Av-mRFP. Histogram was adjusted for visualization purposes. Embryos are oriented top-posterior, bottom-anterior.

**Movie S3**. Live imaging of *frs-MS2*

(left) WT, NC11-NC14; (middle) haploid, NC11-NC15; (right) short-cycle diploid NC11-NC13. *MS2* signal is shown in green. Nuclei are marked with His2Av-mRFP. Histogram was adjusted for visualization purposes.

## References

1. Newport J, Kirschner M (1982) A major developmental transition in early xenopus embryos: I. characterization and timing of cellular changes at the midblastula stage. Cell 30(3):675–686.

2. Newport J, Kirschner M (1982) A major developmental transition in early xenopus embryos: II. control of the onset of transcription. Cell 30(3):687–696.

3. Harrison MM, Eisen MB (2015) Transcriptional Activation of the Zygotic Genome in Drosophila. Current Topics in Developmental Biology doi:10.1016/bs.ctdb.2015.07.028.

4. Blythe SA, Wieschaus EF (2015) Coordinating Cell Cycle Remodeling with Transcriptional Activation at the Drosophila MBT. Current Topics in Developmental Biology, pp 113–148.

5. Yuan K, Seller CA, Shermoen AW, O’Farrell PH (2016) Timing the Drosophila Mid-Blastula Transition: A Cell Cycle-Centered View. Trends Genet 32(8):496–507.

6. Jukam D, Shariati SAM, Skotheim JM (2017) Zygotic Genome Activation in Vertebrates. Dev Cell 42(4):316–332.

7. Liu B, Grosshans J (2017) Link of Zygotic Genome Activation and Cell Cycle Control. Methods in Molecular Biology, pp 11–30.

8. Lefebvre F, Lécuyer É (2018) Flying the RNA Nest: Drosophila Reveals Novel Insights into the Transcriptome Dynamics of Early Development. J Dev Biol 6(1):5.

9. Vastenhouw NL, Cao WX, Lipshitz HD (2019) The maternal-to-zygotic transition revisited. Development. doi:10.1242/dev.161471.

10. Schulz KN, Harrison MM (2019) Mechanisms regulating zygotic genome activation. Nat Rev Genet 20(4):221–234.

11. Edgar BA, Kiehle CP, Schubiger G (1986) Cell cycle control by the nucleo-cytoplasmic ratio in early Drosophila development. Cell 44(2):365–372.

12. Almouzni G, Wolffe AP (1995) Constraints on transcriptional activator function contribute to transcriptional quiescence during early Xenopus embryogenesis. EMBO J 14(8):1752–1765.

13. Lee DR, Lee JE, Yoon HS, Roh S Il, Kim MK (2001) Compaction in preimplantation mouse embryos is regulated by a cytoplasmic regulatory factor that alters between 1- and 2-cell stages in a concentration-dependent manner. J Exp Zool. doi:10.1002/jez.1036.

14. Dekens MPS, Pelegri FJ, Maischein HM, Nüsslein-Volhard C (2003) The maternal-effect gene futile cycle is essential for pronuclear congression and mitotic spindle assembly in the zebrafish zygote. Development. doi:10.1242/dev.00606.

15. Lu X, Li JM, Elemento O, Tavazoie S, Wieschaus EF (2009) Coupling of zygotic transcription to mitotic control at the Drosophila mid-blastula transition. Development 136(12):2101–2110.

16. Jevtić P, Levy DL (2015) Nuclear Size Scaling during Xenopus Early Development Contributes to Midblastula Transition Timing. Curr Biol 25(1):45–52.

17. Chen H, Einstein LC, Little SC, Good MC (2019) Spatiotemporal Patterning of Zygotic Genome Activation in a Model Vertebrate Embryo. Dev Cell 49(6):852–866.e7.

18. Chan SH, et al. (2019) Brd4 and P300 Confer Transcriptional Competency during Zygotic Genome Activation. Dev Cell 49(6):867–881.e8.

19. Ferree PL, Deneke VE, Di Talia S (2016) Measuring time during early embryonic development. Semin Cell Dev Biol 55:80–88.

20. Shermoen AW, O’Farrell PH (1991) Progression of the cell cycle through mitosis leads to abortion of nascent transcripts. Cell 67(2):303–310.

21. Rothe M, Pehl M, Taubert H, Jäckle H (1992) Loss of gene function through rapid mitotic cycles in the Drosophila embryo. Nature 359(6391):156–159.

22. Edgar BA, Schubiger G (1986) Parameters controlling transcriptional activation during early drosophila development. Cell. doi:10.1016/0092-8674(86)90009-7.

23. Kimelman D, Kirschner M, Scherson T (1987) The events of the midblastula transition in Xenopus are regulated by changes in the cell cycle. Cell 48(3):399–407.

24. Collart C, Allen GE, Bradshaw CR, Smith JC, Zegerman P (2013) Titration of Four Replication Factors Is Essential for the Xenopus laevis Midblastula Transition. Science (80-) 341(6148):893–896.

25. Strong I, Yuan K, O’Farrell PH (2017) Interphase-arrested *Drosophila* embryos initiate Mid-Blastula Transition at a low nuclear-cytoplasmic ratio. bioRxiv:143719.

26. Newport J, Dasso M (1989) On the coupling between DNA replication and mitosis. J Cell Sci 1989(Supplement 12):149–160.

27. Clute P, Masui Y (1995) Regulation of the Appearance of Division Asynchrony and Microtubule-Dependent Chromosome Cycles in Xenopus laevis Embryos. Dev Biol 171(2):273–285.

28. Müller F, Lakatos L, Dantonel J-C, Strähle U, Tora L (2001) TBP is not universally required for zygotic RNA polymerase II transcription in zebrafish. Curr Biol 11(4):282–287.

29. Hadzhiev Y, et al. (2019) A cell cycle-coordinated Polymerase II transcription compartment encompasses gene expression before global genome activation. Nat Commun 10(1):691.

30. McKnight SL, Miller Jr. OL (1976) Ultrastructural patterns of RNA synthesis during early embryogenesis of Drosophila melanogaster. Cell 8(2):305–319.

31. De Renzis S, Elemento O, Tavazoie S, Wieschaus EF (2007) Unmasking Activation of the Zygotic Genome Using Chromosomal Deletions in the Drosophila Embryo. PLoS Biol 5(5):e117.

32. Heyn P, et al. (2014) The Earliest Transcribed Zygotic Genes Are Short, Newly Evolved, and Different across Species. Cell Rep 6(2):285–292.

33. Kwasnieski JC, Orr-Weaver TL, Bartel DP (2019) Early genome activation in Drosophila is extensive with an initial tendency for aborted transcripts and retained introns. Genome Res 29(7):1188–1197.

34. Edgar BA, Datar SA (1996) Zygotic degradation of two maternal Cdc25 mRNAs terminates Drosophila’s early cell cycle program. Genes Dev 10(15):1966–1977.

35. Shermoen AW, McCleland ML, O’Farrell PH (2010) Developmental Control of Late Replication and S Phase Length. Curr Biol 20(23):2067–2077.

36. Sung H, Spangenberg S, Vogt N, Großhans J (2013) Number of Nuclear Divisions in the Drosophila Blastoderm Controlled by Onset of Zygotic Transcription. Curr Biol 23(2):133–138.

37. Amodeo AA, Jukam D, Straight AF, Skotheim JM (2015) Histone titration against the genome sets the DNA-to-cytoplasm threshold for the Xenopus midblastula transition. Proc Natl Acad Sci 112(10):E1086–E1095.

38. Jevtić P, Levy DL (2017) Both Nuclear Size and DNA Amount Contribute to Midblastula Transition Timing in Xenopus laevis. Sci Rep. doi:10.1038/s41598-017-08243-z.

39. Pritchard DK, Schubiger G (1996) Activation of transcription in Drosophila embryos is a gradual process mediated by the nucleocytoplasmic ratio. Genes Dev 10(9):1131–1142.

40. Veenstra GJC, Destrée OHJ, Wolffe AP (1999) Translation of Maternal TATA-Binding Protein mRNA Potentiates Basal but Not Activated Transcription in Xenopus Embryos at the Midblastula Transition. Mol Cell Biol. doi:10.1128/mcb.19.12.7972.

41. Guven-Ozkan T, Nishi Y, Robertson SM, Lin R (2008) Global Transcriptional Repression in C. elegans Germline Precursors by Regulated Sequestration of TAF-4. Cell. doi:10.1016/j.cell.2008.07.040.

42. Yamada S, et al. (2019) The Drosophila Pioneer Factor Zelda Modulates the Nuclear Microenvironment of a Dorsal Target Enhancer to Potentiate Transcriptional Output. Curr Biol 29(8):1387–1393.e5.

43. Bertrand E, et al. (1998) Localization of ASH1 mRNA particles in living yeast. Mol Cell. doi:10.1016/S1097-2765(00)80143-4.

44. Forrest KM, Gavis ER (2003) Live Imaging of Endogenous RNA Reveals a Diffusion and Entrapment Mechanism for nanos mRNA Localization in Drosophila. Curr Biol 13(14):1159–1168.

45. Golding I, Paulsson J, Zawilski SM, Cox EC (2005) Real-Time Kinetics of Gene Activity in Individual Bacteria. Cell 123(6):1025–1036.

46. Larson DR, Zenklusen D, Wu B, Chao JA, Singer RH (2011) Real-Time Observation of Transcription Initiation and Elongation on an Endogenous Yeast Gene. Science (80-) 332(6028):475–478.

47. Garcia HG, Tikhonov M, Lin A, Gregor T (2013) Quantitative imaging of transcription in living Drosophila embryos links polymerase activity to patterning. Curr Biol. doi:10.1016/j.cub.2013.08.054.

48. Fukaya T, Lim B, Levine M (2016) Enhancer Control of Transcriptional Bursting. Cell 166(2):358–368.

49. Loppin B, Docquier M, Bonneton F, Couble P (2000) The maternal effect mutation sesame affects the formation of the male pronucleus in Drosophila melanogaster. Dev Biol. doi:10.1006/dbio.2000.9718.

50. Loppin B, Berger F, Couble P (2001) The Drosophila maternal gene sésame is required for sperm chromatin remodeling at fertilization. Chromosoma 110(6):430–440.

51. Loppin B, et al. (2005) The histone H3.3 chaperone HIRA is essential for chromatin assembly in the male pronucleus. Nature. doi:10.1038/nature04059.

52. Di Talia S, et al. (2013) Posttranslational Control of Cdc25 Degradation Terminates Drosophila’s Early Cell-Cycle Program. Curr Biol 23(2):127–132.

53. Farrell JA, O’Farrell PH (2013) Mechanism and Regulation of Cdc25/Twine Protein Destruction in Embryonic Cell-Cycle Remodeling. Curr Biol 23(2):118–126.

54. Blythe SA, Wieschaus EF (2016) Establishment and maintenance of heritable chromatin structure during early Drosophila embryogenesis. Elife 5. doi:10.7554/eLife.20148.

55. Shindo Y, Amodeo AA (2019) Dynamics of Free and Chromatin-Bound Histone H3 during Early Embryogenesis. Curr Biol 29(2):359–366.e4.

56. Fogarty P, Kalpin RF, Sullivan W (1994) The Drosophila maternal-effect mutation grapes causes a metaphase arrest at nuclear cycle 13. Development 120(8):2131–2142.

57. Sibon OCM, Stevenson VA, Theurkauf WE (1997) DNA-replication checkpoint control at the Drosophila midblastula transition. Nature 388(6637):93–97.

58. Blythe SA, Wieschaus EF (2015) Zygotic Genome Activation Triggers the DNA Replication Checkpoint at the Midblastula Transition. Cell 160(6):1169–1181.

59. Foo SM, et al. (2014) Zelda Potentiates Morphogen Activity by Increasing Chromatin Accessibility. Curr Biol 24(12):1341–1346.

60. Ferraro T, et al. (2016) Transcriptional Memory in the Drosophila Embryo. Curr Biol 26(2):212–218.

61. Lott SE, et al. (2011) Noncanonical compensation of zygotic X transcription in early Drosophila melanogaster development revealed through single-embryo RNA-Seq. PLoS Biol. doi:10.1371/journal.pbio.1000590.

62. Chari S, Wilky H, Govindan J, Amodeo AA (2019) Histone concentration regulates the cell cycle and transcription in early development. Development 146(19):dev177402.

63. Djabrayan NJV, et al. (2019) Metabolic Regulation of Developmental Cell Cycles and Zygotic Transcription. Curr Biol 29(7):1193–1198.e5.

## SI References

1. Fukaya T, Lim B, Levine M (2016) Enhancer Control of Transcriptional Bursting. Cell 166(2):358–368.

2. Djabrayan NJV, et al. (2019) Metabolic Regulation of Developmental Cell Cycles and Zygotic Transcription. Curr Biol 29(7):1193–1198.e5.

3. Gratz SJ, et al. (2014) Highly Specific and Efficient CRISPR/Cas9-Catalyzed Homology-Directed Repair in Drosophila. Genetics 196(4):961–971.

4. Lim B, Fukaya T, Heist T, Levine M (2018) Temporal dynamics of pair-rule stripes in living Drosophila embryos. Proc Natl Acad Sci U S A 115(33):8376–8381.

